# Epithelial stratification shapes infection dynamics

**DOI:** 10.1101/231985

**Authors:** Carmen Lía Murall, Robert Jackson, Ingeborg Zehbe, Nathalie Boulle, Michel Segondy, Samuel Alizon

## Abstract

Infections of stratified epithelia collectively represent a large burden on global health. Experimental models provide a means to understand how the cell dynamics themselves influence the outcomes of these infections. Mathematical approaches are needed to improve quantification and theoretical advancement of these complex systems. Here, we develop a general ecology-inspired model for stratified epithelial dynamics, which allows us to simulate infections and to estimate parameters that are difficult to measure with organotypic cell cultures. To explore how epithelial cell dynamics affect infection dynamics, we focus on two contrasting pathogens of the cervicovaginal epithelium: *Chlamydia trachomatis* and Human papillomaviruses. We find that key infection symptoms stem from differential interactions with the layers, while clearance and pathogen burden are bottom-up processes. Cell protective responses to infections (e.g. increased cell proliferation) generally lowered pathogen load but there were specific effects based on infection strategies. These generic responses by the epithelium, then, will have varying results depending on the pathogen’s infection strategy. Our modeling approach opens new perspectives for 3D tissue culture experimental systems of infections and, more generally, for developing and testing hypotheses related to infections of stratified epithelia.

## I. INTRODUCTION

Localized (non-systemic) infections of stratified epithelia are a broad class of infections that collectively represent a major burden on global public health systems. For instance, skin diseases, a majority of which are caused by infections, are the 18^th^ leading cause of health burden worldwide (in disability-adjusted life years, DALY rank) and bacterial skin infections alone cause 66,500 annual deaths (from the 2010 Global Burden of Disease study [1]). Furthermore, many sexually transmitted infections (STIs) are epithelial and are globally on the rise [2]. In 2016, the World Health Organization (WHO) estimated that four curable STIs (*Chlamydia trachomatis*, *Neisseria gonorrhoeae*, syphilis and *Trichomonas vaginalis*) cause 357 million new infections worldwide every year. They also estimate that 417 million people carry Herpes simplex virus type 2 (HSV-2) and 291 million women have ano-genital infections by Human papillomaviruses worldwide (HPVs) [3], however, due to the asymptomatic nature of many of these infections the global prevalence is likely higher. In addition, *Mycoplasma genitalium* and *N. gonorrhoeae* are increasingly becoming resistant to antibiotics [4], thus causing rising morbidity, mortality and costs [2]. Facing these challenges requires a better understanding of epithelial dynamics.

While stratified epithelia are often the first line of defense against infections [5], their cells are often the primary resource for many viruses or bacteria. This is why understanding epithelial life-cycles, signaling, and dynamics is an active line of research [6]. Epithelial infections are very heterogeneous in their outcomes, from short sub-clinical acute infections to chronic pathologies [1]. Our hypothesis is that the epithelial stratified structure is one of the keys to understanding these patterns. Though experimental and clinical methods used for studying these infections are increasingly more quantitative (e.g. flow cytometry or -omics technologies), theoretical frameworks for understanding infection properties and dynamics in stratified epithelia are lacking since most models consider infections of monolayers or the blood. Here, we build on the analogy between a host and an ecological system [7, 8] to investigate how the stratification of the epithelium drives infection dynamics, with a particular focus on the cervicovaginal epithelium and STIs.

Localized STIs of the cervicovaginal mucosa are involved in a range of health concerns, such as decreasing fertility [3, 9–11] or carcinogenesis [12]. Studying the cervical epithelium has greatly helped improve women’s health [13] and histological studies of cervical infections have characterized both healthy and diseased cells. The ectocervix is a non-keratinized stratified epithelium that acts as an important barrier to prevent infections from entering the upper part of the female genital tract and affecting fertility. The tight packing of the epithelial cells and their migration to the surface are believed to prevent bacteria and viruses from reaching the dermis [14]. Furthermore, the continual production of surface mucus is thought to aid in trapping and removing invaders [15]. Studying these processes using tractable experimental systems has been a challenge given the complexity of recreating stratified epithelia with realistic features, but this is changing rapidly [16]. Mathematical modeling can aid this experimental work by helping to estimate parameters such as changes in cell migration or mucus production rate.

The vast majority of mathematical models of STI within-host dynamics focus on HIV (for a review, see [17]) but some investigate pathogens that only (or mainly) target epithelia such as Chlamydia [18–22], HPV [23–26], Epstein-Barr Virus (EBV) [27, 28] or HSV [29]. A common feature of these models is that they focus on the pathogen and the associated immune response somehow at the expenses of the epithelium itself. As a consequence, with few exceptions (e.g. [25]), they assume that the population of cells infected by the pathogen is homogeneous and not structured. We take an ecological approach to model the stratified epithelium to investigate the effect of the structure of the life cycle of the host cells on infection dynamics. The analogy between ecological systems and pathogen-host interactions is not new (e.g. [7]), but it is becoming increasingly common and has underlaid successful quantitative tools for understanding viral kinetics [17, 30] and drug resistance [31].

From an ecological perspective, the stratified epithelial structure can be viewed as having stages or age structure (herein called ‘*stage-structure*’), meaning the full lifecycle of an ‘individual’ is divided up into stages or ages. Therefore, populations of one phenotype (or age) give rise to another in a successive fashion. Ecological populations with stage-structure have been shown to have rich dynamics that impact the entire ecological system [32–34] and generally oversimplifying resources is known to potentially lead to incorrect predictions [35]. Similar importance has been shown in epidemiology, where age structure of host populations and host heterogeneity are key determinants of infection spread [36–38]. At the within-host level, studies have shown the importance of the target cells age. For instance, by combining mathematical models with experimental data, Mideo *et al.* showed that differences between *Plasmodium chabaudi* strains could be most parsimoniously explained by their different affinity for erythrocytes of different ages, as well as differences in erythropoiesis [39]. Target cell heterogeneity has also been put forward to explain the HIV co-receptor switch [40]. Our hypothesis, then, is that the stratified structure is key to understanding various epithelial infections. While we pursue this analogy, we insist that stratified epithelia exhibit features that differ from traditional populations. For instance, differentiated keratinocytes (or ‘adults’) do not reproduce to make stem cells (or ‘juveniles’) the way free-living species do. Additionally, the epithelium self-regulates its dynamics as a means to maintain homeostasis (e.g. changing differentiation and shedding). This, therefore, calls for a system-specific approach.

Having a framework for epithelial dynamics allows us to simulate infections. For this, we chose two extremely prevalent STIs with very different biological features: Human papillomaviruses (low-risk, LR, and high-risk, HR) and *C. trachomatis* bacteria. In the United States alone, more than 1.5 million cases of *C. trachomatis* were reported to the Center for Disease Control (CDC) in 2015 and HPVs are the most common STI in the country [41]. While both Chlamydia and HPV replicate intra-cellularly, these two infections exhibit contrasting strategies for infecting the squamous epithelium: HPVs cause non-lytic basal-up infections, whereas Chlamydia infections are from the surface-down and are lytic. As mentioned earlier, there are some previous mathematical models of both HPV and Chlamydia, which we can build on [18–26]. More importantly, the biology of these two pathogens are well studied, with their life-histories (HPVs [42] and Chlamydia [43]) well characterized, at least, more so than other epithelial infections. This provided us with peer-reviewed parameter estimates and assumptions for our model. Consequently, we needed to make few assumptions about the biology and could compare our results to previous mathematical models without age-structure of the epithelium. Finally, in order to maintain focus on the epithelium, we used a simple model for the immune response, as in earlier studies (e.g. on HSV, [29]).

We address to what extent epithelium dynamics affect infection dynamics and as a result determine infection outcomes. First, we introduce a general epithelium model, which we calibrate using existing data, as well as original cell culture data from a spontaneously immortalized human cell line (NIKS) [44]. With this data we infer parameters that are difficult to measure, such as the probability of symmetric cell divisions. We then ‘infect’ this epithelial model with LR and HR HPVs and Chlamydia to investigate how protective measures by the epithelium affect infection load and duration, while identifying the parameters that control key infection traits. We find that epithelium stratification plays a key role in the dynamics and outcomes of these infections.

## RESULTS

### Uninfected epithelial dynamics

Our model abstracts the stratified epithelium into four phenotypically distinct populations, relevant to clinical and experimental models of the epithelium: stem-like cells in the basal and parabasal layers, and differentiated cells in the mid and surface cells (Figure 1 and equation system 1). These phenotypes can be identified experimentally using immunofluorescence techniques that target genes or proteins expressed differentially as the cells mature and move up the epithelial column. The model includes 7 parameters, of which 4 are inferred from autoradiographic experiments done in 1970 [45] and one, *N_b_*, is a scaling parameter describing the surface of the basal monolayer considered. The two remaining free parameters capture the difference in symmetric divisions probabilities by the basal and parabasal cells (Δ_*p*_ and Δ_*q*_). All the parameters are listed in Table I.

**FIG. 1.**
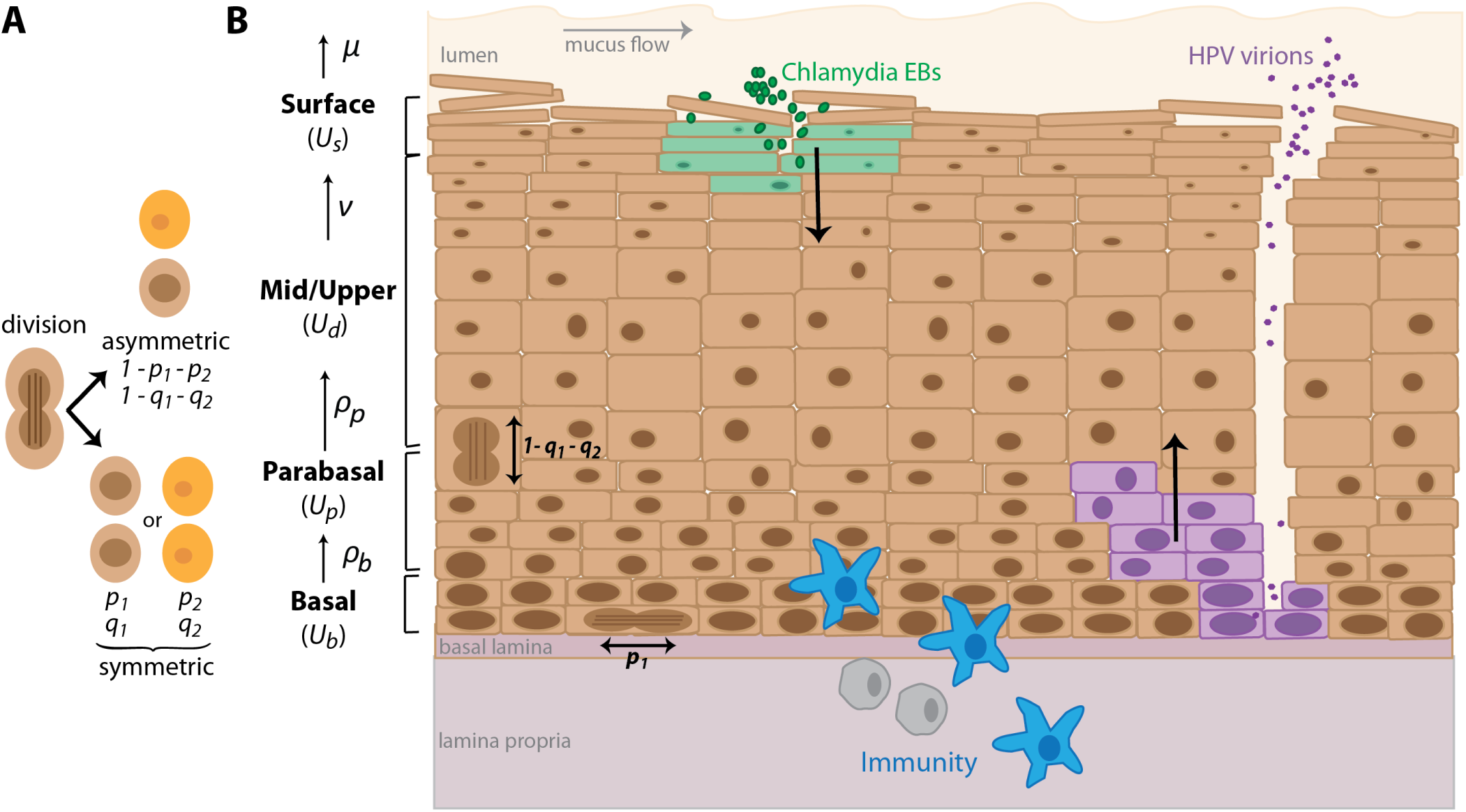
Dynamics in the stratified epithelium. A) Basal and parabasal cells can divide either asymmetrically (1 − *p*_1_ − *p*_2_ and 1 − *q*_1_ − *q*_2_ respectively) or symmetrically, which result in two daughter cells of either the same (*p*_1_ or *q*_1_) or different phenotype as the mother cell (*p*_2_ or *q*_2_). B) The squamous epithelium is abstracted into a basal, a parabasal, a mid-upper and a surface layer. Proliferation (*ρ*) and maturation (*ν*) rates determine the movement of cells up the layers. Cells die and are shed (*μ*). *Chlamydia trachomatis* (in green) infects the most superficial live cells through the mucus and underneath shedding dead cells. Once inside a cell, the Elementary Bodies (EBs) change into Reticulate Bodies (RBs), which go through several rounds of replication, and then change back into EBs that are released upon cell death. *Human papillomaviruses* (in purple) must infect basal cells to establish an infection, thus usually requiring a micro-abrasion. The virus is non-lytic and replicates in host cells as they follow their natural life-cycle up the epithelium column. Progeny virions are released once the cell dies at the surface. Immune cells (in blue) enter the epithelium from the basal layer.

**TABLE I.** Parameter descriptions for epithelial model, default values, biologically realistic ranges and estimated values. Parameter values that were chosen for the results to be biologically consistent are labeled as ‘calibrated’. The values ‘fixed*’ and ‘estimated*’ were derived using data (see Supplementary Information). The data from NIKS cell cultures, which are a common cell-line used to model these systems, but are not identical to *in vivo* cells in the cervicovaginal squamous epithelium (e.g. the latter cannot form keratinized layers).

|  |  | Default | Range | Ref |
| --- | --- | --- | --- | --- |
| *a. in vivo estimates (literature)* |  |  |  |  |
| $N_b$ | Total number of basal cells | $10^3$ | $[10^2 ; 10^5]$ | fixed |
| $\rho_b$ | Basal cell replication rate ($\text{day}^{-1}$) | 0.03 | $[0.03 ; 0.07]$ | [45] |
| $\rho_p$ | Parabasal cell replication rate ($\text{day}^{-1}$) | 0.39 | $[0.2 ; 1]$ | [45] |
| $\Delta p$ | Difference of symmetric divisions (basal to parabasal) | -0.012 | $[-1 ; 1]$ | calibrated |
| $\Delta q$ | Difference of symmetric divisions (parabasal to differentiated) | -0.012 | $[-1 ; 1]$ | calibrated |
| $\nu$ | Keratinocyte migration rate (mid/upper to surface layer, $\text{day}^{-1}$) | 0.4 | $[0.2 ; 1]$ | [45] |
| $\mu$ | Keratinocyte natural death rate ($\text{cell}^{-1}.\text{day}^{-1}$) | 0.67 | $[0.2 ; 1]$ | [45] |
| *b. culture estimates (data-derived)* |  |  |  |  |
| $N_b$ | Basal cell carrying capacity per FOV | 42 | $[- ; -]$ | fixed* |
| $\rho_b$ | Basal cell replication rate ($\text{day}^{-1}$) | 0.058 | $[0.06 ; 0.062]$ | estimated |
| $\rho_p$ | Parabasal cell replication rate ($\text{day}^{-1}$) | 0.0077 | $[- ; -]$ | estimated* |
| $\Delta p$ | Difference of symmetric divisions (basal to parabasal) | $\approx 0$ | $[0 ; 0.018]$ | estimated |
| $\Delta q$ | Difference of symmetric divisions (parabasal to differentiated) | -0.99 | $[-1 ; -0.96]$ | estimated |
| $\nu$ | Keratinocyte migration rate (mid/upper to surface layer, $\text{day}^{-1}$) | 0.20 | $[0.16 ; 0.23]$ | estimated |
| $\mu$ | Keratinocyte natural death rate ($\text{cell}^{-1}.\text{day}^{-1}$) | $3.4 \times 10^{-8}$ | $[0.010 ; 0.015]$ | estimated |

The parameters for which we have less information are related to the probability of cells dividing symmetrically (that is for a basal stem cell to produce two daughter basal stem cells or, conversely, two differentiated cells). Existing data suggests that such divisions are rare [46, 47]. This is further reinforced from our estimation of epithelium thickness. Histological studies estimate 26 to 28 cell layers in the vaginal epithelium depending on the stage of the menstrual cycle [48] and *in vivo* studies of the cervical epithelium count 16 to 17 layers [49]. To achieve comparable values, and assuming that the ranges of the other parameters are biologically plausible, we find that symmetric divisions must be rare. Calibrating Δ_*q*_ ≈ −0.012 gives an epithelium ‘thickness’ of 17 layers, i.e. 17*N_b_*. Analytical results shown in Supporting Information revealed the need for some degree of symmetric division biased towards producing differentiated cells (Δ_*q*_ < 0). Furthermore, if we assume each layer of parabasal cells has the same number of basal cells and that the differentiated cells are half the number of cells per layer (because they are twice the size [50]), then 17*N_b_* corresponds to 26 layers. Finally, we found that the mid layers, that is the differentiated (*U_d_*) and parabasal layers (*U_p_*), are larger than the basal and superficial (*U_s_*) layers (this can be seen in the pre-infection period in Figures 3 and 4).

In order to obtain experimentally relevant parameter estimates, we used our model and the known parameters as priors to estimate values using original data from raft cultures of NIKS cells. Figure 2A and B show an example of epithelial growth into stratified form. Figure 2C shows the dynamics of the number of basal and suprabasal (non-keratinized and keratinized) cells, along with the inferred dynamics from the model. In this data of a period of growth (from single layer to stratified), the symmetric divisions were inferred to be negligible in the basal layer but important in the parabasal layers (Table I). The inverse relationship between *ρ_p_* and Δ_*q*_ implies that since the data constrained *ρ_p_* to be low (see Supporting Information) then the high estimated value for Δ_*q*_ suggests that nearly all divisions produce two differentiated cells (Table I). This, along with the higher than default basal replication rates Table I), is consistent with a growth phase of an epithelium up to homeostasis.

**FIG. 2.**
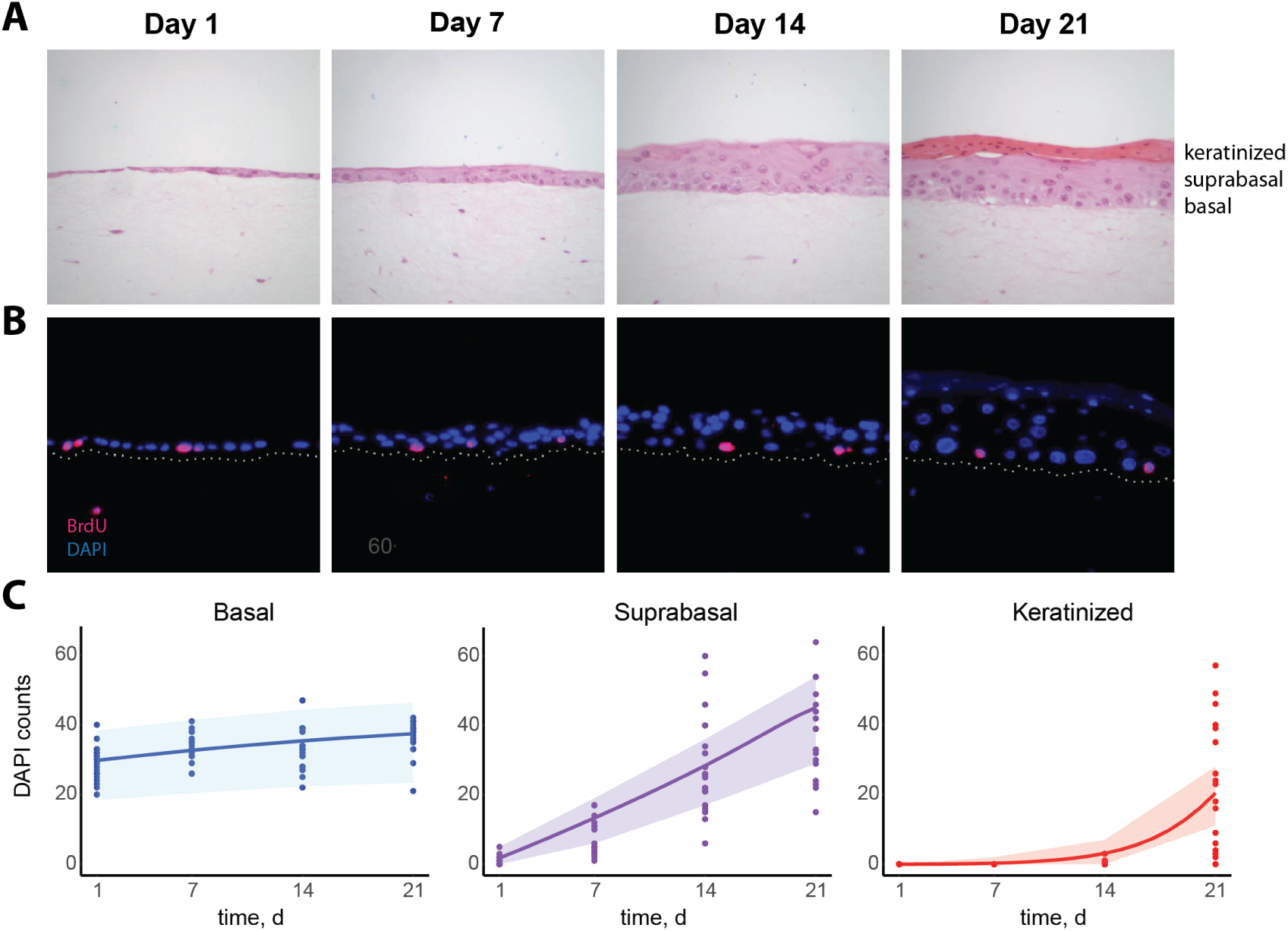
Epithelial cell growth in 3D raft cultures. A) NIKS cells grown from a single layer over a period of three weeks. Dark pink layer in week 3 are cornified cells that accumulate on the surface. B) Immunoflourescence staining, DAPI (blue) and BrdU (pink); white dots are added basal lines. C) Data of NIKS growth over time with model fitting. Shading corresponds to 95% prediction interval, assuming the data follows a Poisson distribution.

We performed a sensitivity analysis to explore the general behaviour of the model and identify the parameters that have the largest effect on homeostasis (Table II). This showed that the total number of cells in the layers above the basal layer is mostly governed by the basal cell proliferation rate (*ρ_b_*) and the difference of symmetric divisions producing parabasal cells (Δ_*p*_). Additionally, the time for the system to reach homeostasis (which is important for repairing damaged tissues) depends on the proliferation rate of the parabasal cells (*ρ_p_*; Supporting Information). Indeed, this is also found from fitting the data and simulated data, where the replication rate, *ρ_p_*, or the symmetric divisions of the parabasal cells, Δ_*q*_, are significantly higher as homeostasis is reached faster (Table I and not shown).

**TABLE II.**
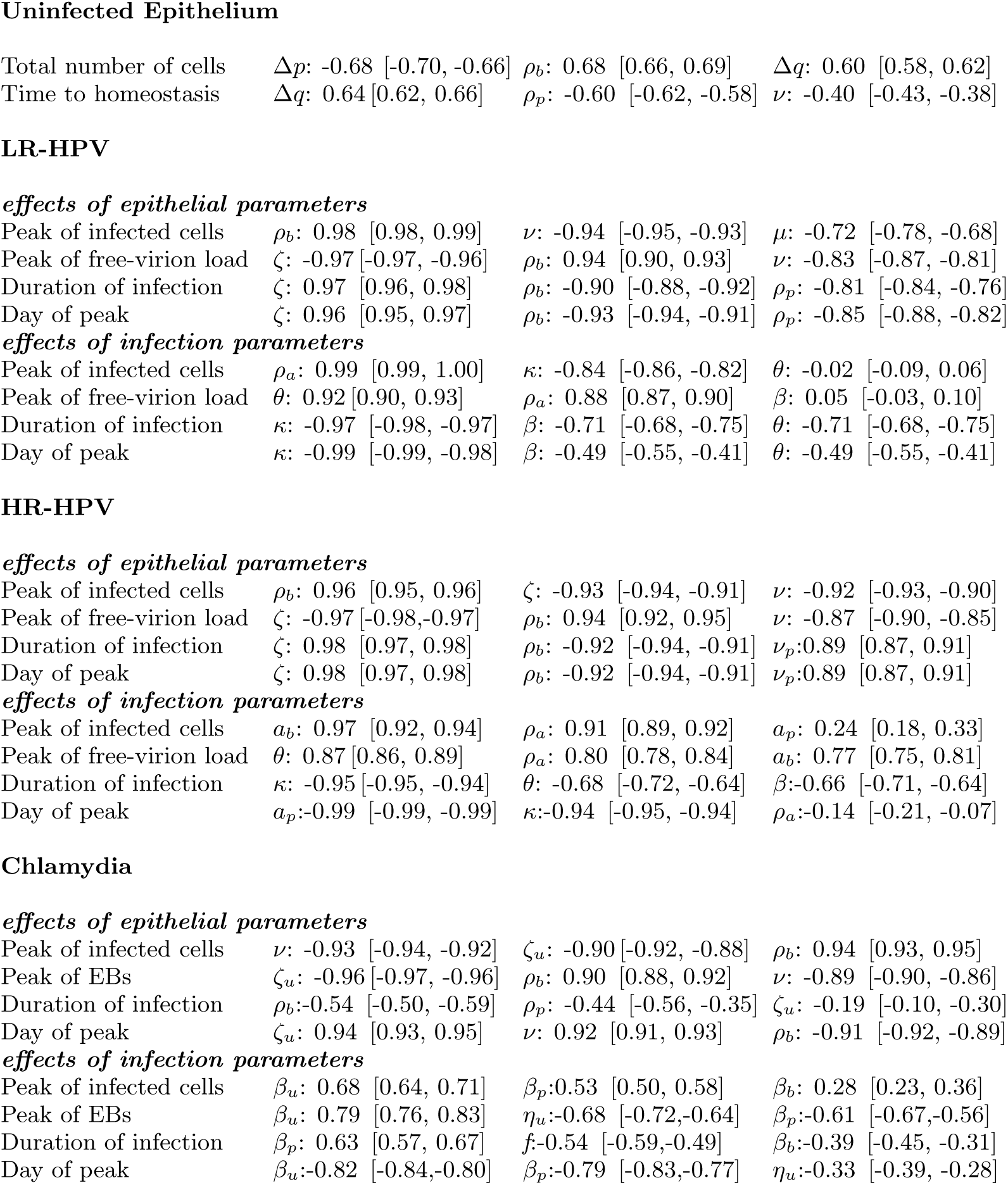
Sensitivity analyses of key infection properties. For each pathogen, we show the three most important parameters with the associated PRCC and its 95% confidence interval. Notations for parameter values are in Table I. Peak of infected cells is a measure of the size of the infection, peak of free-virion (or EB) load is how much progeny is released for re-seeding the infection or transmission, and day of peak is a measure for how quickly the infection grows. For effects on protection by epithelial parameters we tested: *ν*, *μ*, *ρ_b_*, *ρ_p_*, *ζ*, and *ζ_u_*

Uninfected Epithelium
|  |  |  |  |
| --- | --- | --- | --- |
| Total number of cells | $\Delta p$: -0.68 [-0.70, -0.66] | $\rho_b$: 0.68 [0.66, 0.69] | $\Delta q$: 0.60 [0.58, 0.62] |
| Time to homeostasis | $\Delta q$: 0.64 [0.62, 0.66] | $\rho_p$: -0.60 [-0.62, -0.58] | $\nu$: -0.40 [-0.43, -0.38] |
| *effects of epithelial parameters* |  |  |  |
| Peak of infected cells | $\rho_b$: 0.98 [0.98, 0.99] | $\nu$: -0.94 [-0.95, -0.93] | $\mu$: -0.72 [-0.78, -0.68] |
| Peak of free-virion load | $\zeta$: -0.97 [-0.97, -0.96] | $\rho_b$: 0.94 [0.90, 0.93] | $\nu$: -0.83 [-0.87, -0.81] |
| Duration of infection | $\zeta$: 0.97 [0.96, 0.98] | $\rho_b$: -0.90 [-0.88, -0.92] | $\rho_p$: -0.81 [-0.84, -0.76] |
| Day of peak | $\zeta$: 0.96 [0.95, 0.97] | $\rho_b$: -0.93 [-0.94, -0.91] | $\rho_p$: -0.85 [-0.88, -0.82] |
| *effects of infection parameters* |  |  |  |
| Peak of infected cells | $\rho_a$: 0.99 [0.99, 1.00] | $\kappa$: -0.84 [-0.86, -0.82] | $\theta$: -0.02 [-0.09, 0.06] |
| Peak of free-virion load | $\theta$: 0.92 [0.90, 0.93] | $\rho_a$: 0.88 [0.87, 0.90] | $\beta$: 0.05 [-0.03, 0.10] |
| Duration of infection | $\kappa$: -0.97 [-0.98, -0.97] | $\beta$: -0.71 [-0.68, -0.75] | $\theta$: -0.71 [-0.68, -0.75] |
| Day of peak | $\kappa$: -0.99 [-0.99, -0.98] | $\beta$: -0.49 [-0.55, -0.41] | $\theta$: -0.49 [-0.55, -0.41] |
| *effects of epithelial parameters* |  |  |  |
| Peak of infected cells | $\rho_b$: 0.96 [0.95, 0.96] | $\zeta$: -0.93 [-0.94, -0.91] | $\nu$: -0.92 [-0.93, -0.90] |
| Peak of free-virion load | $\zeta$: -0.97 [-0.98, -0.97] | $\rho_b$: 0.94 [0.92, 0.95] | $\nu$: -0.87 [-0.90, -0.85] |
| Duration of infection | $\zeta$: 0.98 [0.97, 0.98] | $\rho_b$: -0.92 [-0.94, -0.91] | $\nu_p$: 0.89 [0.87, 0.91] |
| Day of peak | $\zeta$: 0.98 [0.97, 0.98] | $\rho_b$: -0.92 [-0.94, -0.91] | $\nu_p$: 0.89 [0.87, 0.91] |
| *effects of infection parameters* |  |  |  |
| Peak of infected cells | $a_b$: 0.97 [0.92, 0.94] | $\rho_a$: 0.91 [0.89, 0.92] | $a_p$: 0.24 [0.18, 0.33] |
| Peak of free-virion load | $\theta$: 0.87 [0.86, 0.89] | $\rho_a$: 0.80 [0.78, 0.84] | $a_b$: 0.77 [0.75, 0.81] |
| Duration of infection | $\kappa$: -0.95 [-0.95, -0.94] | $\theta$: -0.68 [-0.72, -0.64] | $\beta$: -0.66 [-0.71, -0.64] |
| Day of peak | $a_p$: -0.99 [-0.99, -0.99] | $\kappa$: -0.94 [-0.95, -0.94] | $\rho_a$: -0.14 [-0.21, -0.07] |
| *effects of epithelial parameters* |  |  |  |
| Peak of infected cells | $\nu$: -0.93 [-0.94, -0.92] | $\zeta_u$: -0.90 [-0.92, -0.88] | $\rho_b$: 0.94 [0.93, 0.95] |
| Peak of EBs | $\zeta_u$: -0.96 [-0.97, -0.96] | $\rho_b$: 0.90 [0.88, 0.92] | $\nu$: -0.89 [-0.90, -0.86] |
| Duration of infection | $\rho_b$: -0.54 [-0.50, -0.59] | $\rho_p$: -0.44 [-0.56, -0.35] | $\zeta_u$: -0.19 [-0.10, -0.30] |
| Day of peak | $\zeta_u$: 0.94 [0.93, 0.95] | $\nu$: 0.92 [0.91, 0.93] | $\rho_b$: -0.91 [-0.92, -0.89] |
| *effects of infection parameters* |  |  |  |
| Peak of infected cells | $\beta_u$: 0.68 [0.64, 0.71] | $\beta_p$: 0.53 [0.50, 0.58] | $\beta_b$: 0.28 [0.23, 0.36] |
| Peak of EBs | $\beta_u$: 0.79 [0.76, 0.83] | $\eta_u$: -0.68 [-0.72, -0.64] | $\beta_p$: -0.61 [-0.67, -0.56] |
| Duration of infection | $\beta_p$: 0.63 [0.57, 0.67] | $f$: -0.54 [-0.59, -0.49] | $\beta_b$: -0.39 [-0.45, -0.31] |
| Day of peak | $\beta_u$: -0.82 [-0.84, -0.80] | $\beta_p$: -0.79 [-0.83, -0.77] | $\eta_u$: -0.33 [-0.39, -0.28] |

Having generated and calibrated a model for epithelial dynamics, we could then simulate infections to investigate how stratification affects important properties of the infection.

### Symptoms during infection: disruption of homeostasis

Cervical infections by both Chlamydia and HPVs are heterogeneous in their clinical manifestations. Chlamydia infections can be symptom-less or with clinical manifestations such as cervicitis [51]. The lytic nature of Chlamydia infections reduce the epithelium to lower cell numbers than homeostasis, therefore affecting the integrity of the layers (Figure 3). This is consistent with the cervical erosion observed in Chlamydia-driven cervicitis or in infections by other lytic pathogens such as HSV [52].

**FIG. 3.**
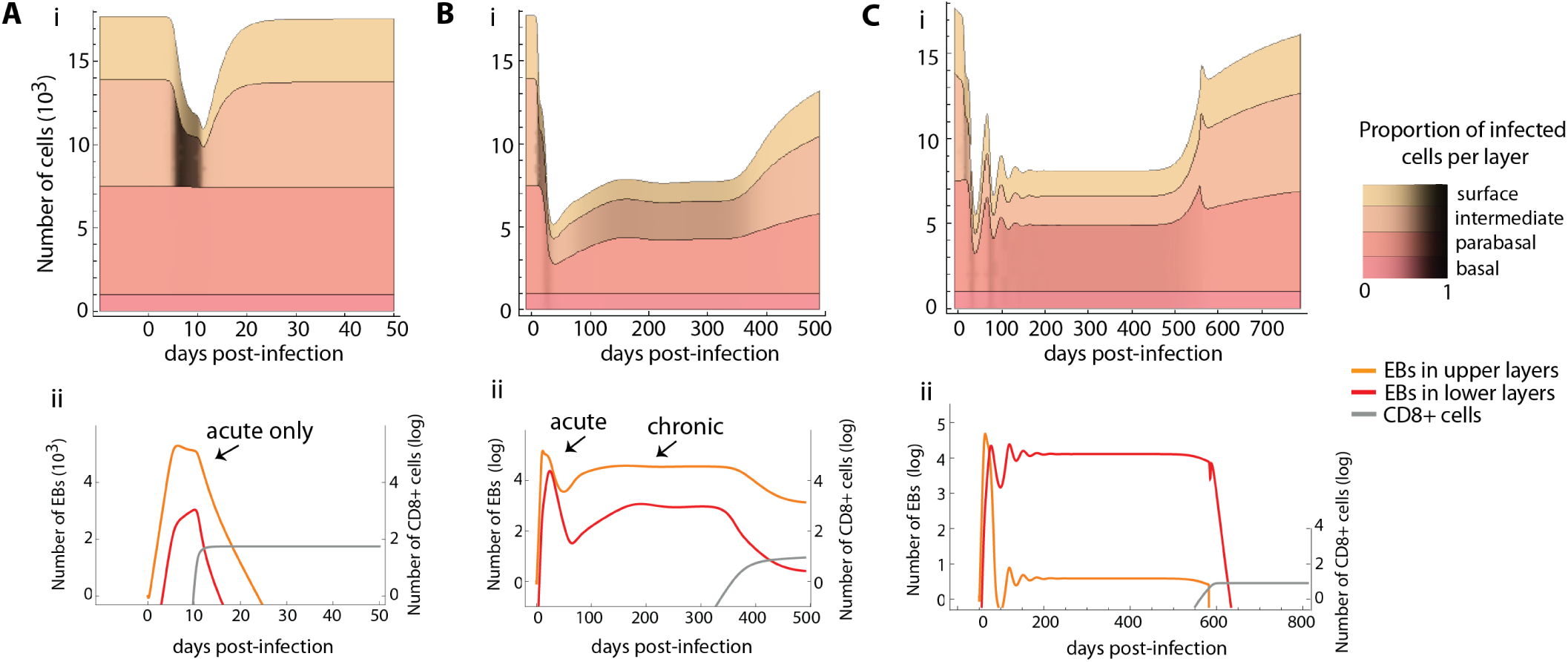
Simulated population dynamics of epithelial cells, immune effectors and bacteria in Chlamydia infections. In A, the immune cell proliferation is rapid, which leads to an acute infection. In B and C, immune cells do not proliferate fast enough to clear the bacteria and the acute phase is followed by oscillations and the establishment of a chronic phase (plateau of EB density). The infection is lytic reducing the thickness of the epithelium (A, B and C) but only chronic infections manage to infect the lower layers (B and C). Parameter values are default (Table I) except for *β_b_* = 8.0 × 10^−7^, *β_p_* = 4.0 × 10^−6^, *β_u_* = 2.0 × 10^−5^, and *ϕ* = 0.0015 in A. In C, all four epithelial protective measures happen together, *ζ_u_*, *rho_b_*, *μ*, and *ν* all rise above default after infection.

LR-HPV types typically generate warts (a substantive overgrowth of cells above homeostasis levels) and HR-HPV types cause flat lesions (often with a thickening of the epithelium) [42]. How differences between HPV types translate into this observed diversity of clinical manifestations in the epithelium is not always clear. The epithelium model allowed us to identify conditions that lead to wart-like symptoms. Assuming that there can be rare events of new virions entering the basal layer (e.g. due to immunosuppression and cytokines loosening epithelial junctions) and that LR types do not drive cell proliferation in lower layers [42], we find that they must either have higher burst sizes (produce more virions per cell) than HR-HPV types or be better at driving differentiated cells back into S-phase in the upper layers (*ρ_a_* and *θ* control the peak of infected cells in Table II). Burst size, *θ*, also controls how quickly the number of infected cells increases, as does the infection rate, *β*. This explains why simulations of LR-HPVs with higher burst sizes are more effective at reaching basal cells, as illustrated by the differences in shading of basal layers between Figure 4A and B.

**FIG. 4.**
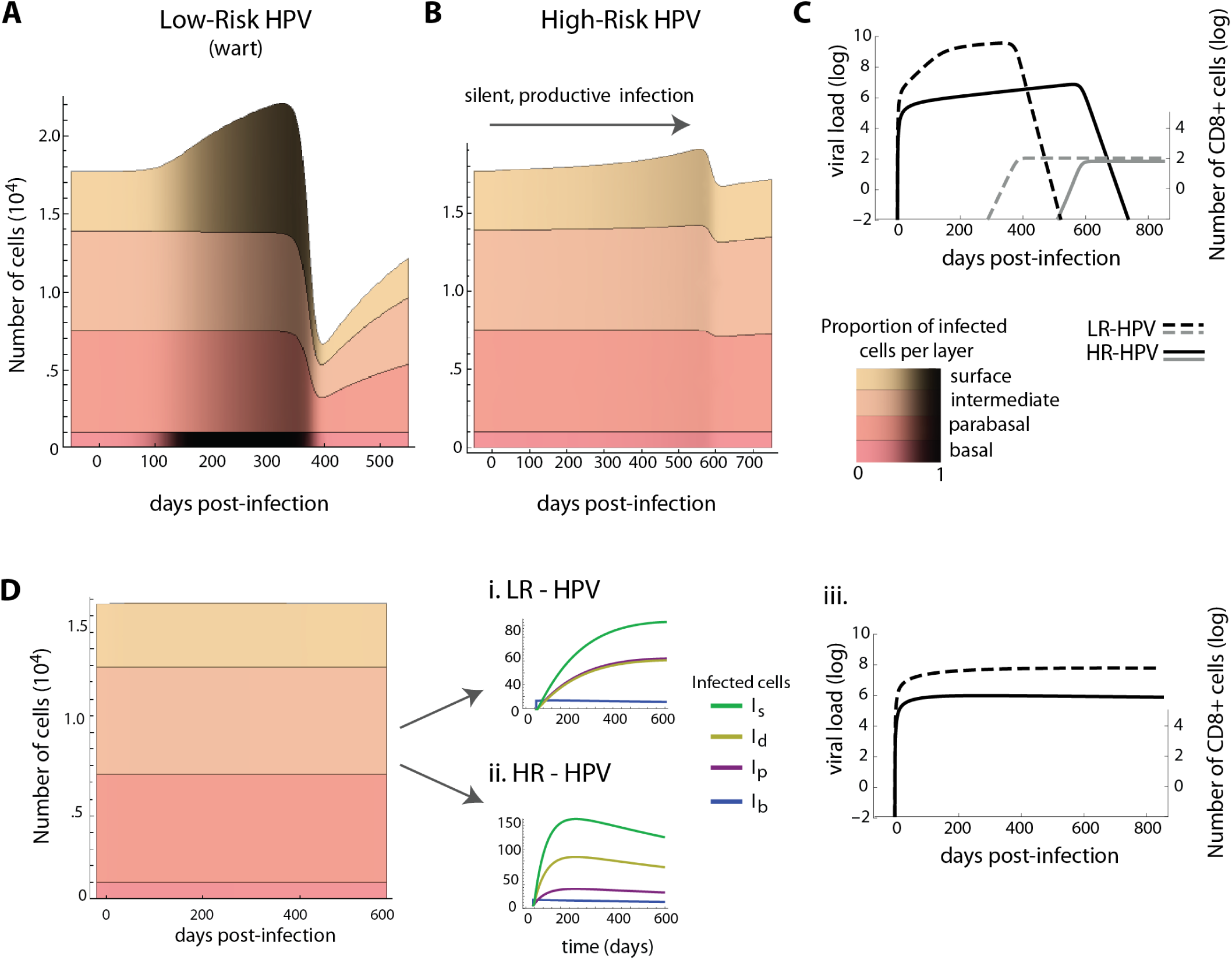
Simulated population dynamics of epithelial cells, immune effectors and free viruses in the case of HPV infections. Epithelial dynamics in a low-risk (LR) HPV infection with a wart-like symptom (A) and a slow growing high-risk (HR) HPV infection that spontaneously regresses (B). The black shading shows the proportion of infected cells in each layer.
(C) Dynamics of virus load (black) and the density of immune effectors (gray) for LR-HPV (dashed line) and HR-HPV (full line) infections. Immune cells start to proliferate upon infection but their number remains below −2 log for several months. (D) Both LR and HR types cause symptom-less infections if the infection is inoculated with few cells and the micro-abrasion repairs quickly. (i, ii, iii) Infections with HR types have more infected cells due to their higher proliferative properties but LR types produce virions. Both infections can last for years as long dynamical transients, if stochasticity or the innate response do not clear them. Parameter values are default (Table I), all infections begin with 10 infected basal cells, and in D *β* decays to zero in 10 days (*b* = −0.5).

HR-HPVs have enhanced E6 and E7 oncoprotein effects in the lower layers [42]. In spite of this increase in epithelial cell division rate their infections are flat, slow growing, and are often clinically indistinguishable from a normal epithelium for many months. For this to occur, we find that the extra proliferation in the basal (*a_b_*) and upper layers (*ρ_a_*) and the type’s burst size (*θ*) must be kept low (Table II and Figure 4B). This implies HR-HPVs would be less ‘productive’ (shed less virions) than LR-HPV during an infection of the same duration (Figure 4C). If HR-HPVs were to have low burst sizes but high oncoprotein-driven proliferation in the lower layers, then their infections would be wart-like (Supporting Information). Thus, to maintain flat lesions the strong HR oncogenes need to be down-regulated. A simulated representation of silent, productive HR-HPV infection is shown in Figure 4B.

Note that for an infection to be sufficiently disruptive to generate a visible manifestation depends on the balance between the size of inoculum (number of cells infected initially) and how quickly the microabrasion closes from repair. For instance, the wart-like overgrowth of cells in Figure 4A can be created either by a small inoculum with slow repair or by a large inoculum and fast repair. When microabrasions close quickly (within a few days) and only a small number of cells are infected initially, both HR and LR types do not cause any visible disruption to homeostasis Figure 4D.

### Infection duration and persistence

For some parameter combinations the kinetics of Chlamydia infections had an acute phase only (Figure 3C), as have been observed in guinea pigs and other models [20]. We obtain this qualitative pattern most readily when the infection rates are the same for all layers or when the lower layers are difficult to infect (for instance due to the reduced permeability down the epithelium column [14]) and the population of immune effectors grows rapidly. From the sensitivity analysis, duration is shortened when increasing the infection rate of the basal layer, *β_b_*, and increasing cell recovery rate (Table II).

We also found an acute phase can be followed by a chronic phase, where a pathogen load stabilizes to a set point value (Figure 3D). How quickly a chronic phase is reached depended on Chlamydia’s infection rates of the various layers. Generally, infection rates had to be low to achieve the chronic phase (because if too high then the bacteria burns through the epithelium and its population crashes). Additionally, if the layers are differentially infected by Chlamydia (i.e. *β_b_* < *β_p_* < *β_u_*), then the chronic phase is reached earlier (see Supporting Information).

In contrast, long-lasting LR and HR HPV infections did not exhibit an acute and a chronic phase in our model. Instead, they persisted either by very long transients or by monotonically reaching an equilibrium (e.g. Figure 4D i and ii). Also, for both HPVs, the immunity killing rate (*κ*) was the most important factor in determining infection duration (Table II). With more antigen in the lower layers to detect, the efficiency of immune killing (*κ*) becomes important for determining duration of infection, speed of growth and size of infected cell accumulation (Table II).

Finally, while Chlamydia and HPVs can cause either acute or chronic infections [53], our model showed that chronicity is achieved through different dynamic mechanisms for each pathogen.

### Protective effect of epithelial dynamics against infection

Upon infection, epithelia exhibit defense mechanisms such as increasing mucus flow, tightening the packing of cells, and increasing proliferation and migration to the surface [54]. We varied epithelial parameters from their homeostasis value to investigate in detail the effect of such mechanisms on various measures of infection using our infection models for HPV and Chlamydia (see Supporting Information equations S1 and S2 and sensitivity analyses in Table II).

We found some mechanisms had similar effect on both HPVs and Chlamydia. First, increasing upward migration of epithelial cells (*ν*) reduced the maximum pathogen load reached during the infection (Table II). Second, mucus trapping (*ζ*) delayed the peak and the duration (although it played a bigger role in decreasing the peak of infection for Chlamydia than for HPV, Table II). And finally, for all infections, increasing basal cell proliferation (*ρ_b_*) scored high in affecting all the infection measures, e.g. size of peak or duration (‘effects of epithelial parameters’ in Table II). However, a pathogen-specific effect was that increasing basal proliferation (*ρ_b_*) of uninfected cells decreases the time to clear HPVs but not Chlamydia. Together, this suggests that epithelial cell features themselves play an important role in infection dynamics and outcomes.

## DISCUSSION

Epithelial infections are a major public health burden and in particular STIs are on the rise causing a worldwide concern [1–3]. Quantitative models, both experimental and mathematical, are essential in developing our understanding of these infections. As for systemic (and virulent) infections such as HIV and HCV, mathematical models have been developed to predict and analyze the kinetics of epithelial infections. Here, we show that to understand these kinetics, it is essential to account for the stratified structure of the epithelium, a property that is absent from most models. We illustrated how such a general framework can be combined with 3D cell culture data to estimate key parameters and how it can generate relevant insights regarding the course of epithelial infections.

### Dynamical implications of ecological features

The rate of basal cell proliferation had a strong effect on the homeostasis of both uninfected and infected epithelia, which suggests an ecological ‘bottom-up controlled’ system [55, 56], analogous to those found in free-living food webs. These bottom-up effects are more apparent if we consider that basal cell replication is strongly determined by the resources that are available in the basal lamina, such as growth factor. While we did not explicitly model the resources of the basal layer (it is implicit in the basal proliferation rates), the growth of the cells in the experimental set-up does depend on concentration and temporo-spatial distribution of growth factors, impacting epithelial thickness and proliferation rates. Therefore, this ecological insight of bottom-up driven systems, could be tested more formally in experimental systems by monitoring resource concentrations.

This bottom-up control is further supported by our finding that accelerating basal cell proliferation, as a response to infection, affected all infection measures (e.g. time of peak, total load, duration). This infection response, then, can have a strong effect on the severity and duration of infections. However, using the same response mechanism might be differentially effective depending on the infection strategy of the pathogen. For instance, we found that increasing cell proliferation did not shorten the infection of Chlamydia. This is probably because proliferation increases the number of uninfected epithelial cells in the upper layers which, for Chlamydia, means more ‘resources’.

Pathogens can have different tropisms for the various cell phenotypes of the stratified epithelium. For instance, EBV more readily infects and replicates in differentiated cells of the upper/mid layers [57], whereas HPV infects the basal layer to establish an infection [42]. We hypothesized that this should impact how effective protective processes (e.g. increased mucus production) of the epithelium are against them. In Chlamydia, where the pathogen infects all cell types equally well, we found that tight packing (i.e. epithelial permeability) mattered to the pathology at the site. The speed at which the epithelium shrank and the stability of the infection system (how quickly it can reach chronic phase) depended on how well the bacteria could access cells down the column. If the bacteria was able to infect the bottom of the column quickly, that led to a population crash due to the lack of resources. On the contrary, and somehow unexpectedly, less epithelial permeability stabilized the infection that then lasted much longer and exhibited a clear chronic phase. This stabilizing effect is also observed in ecological systems when one stage is invulnerable to attack, i.e. a stage refugia [33, 58]. For instance, a parasitic wasp of red scales (a common plant pest) was able to maintain its population, thus function as a biological control, because mature red scales were not vulnerable to attack [33]. Such effects from decreasing permeability (protecting the basal replicative stage) would have implications in the context of treatments that bolster cell adhesion and require testing experimentally.

Considering pathogens with contrasted life-histories allowed us to identify how similar infection outcomes arise. In the case of HR-HPV, persistence was achieved via a slow growth strategy that delays clearance by decreasing the negative dynamical feedback involving the immune system (i.e. faster growth means faster immune detection and clearance). Indeed, HPV types appear to evade, or counteract, these immune responses differently. In particular, viral protein E6 of various HPV types differ in their many cellular binding partners resulting in a variety of effects on host processes [59]. In contrast, the interaction between free form Chlamydia and its infection rates of the various stages drove the chronic phase, but although the activation of the immune response through the same feedback ultimately led to clearance. This feedback affected several infection features. For instance, the difference between HPV-induced genital warts and cervical lesions depended most on the number of virions an infected cell releases upon death (or ‘burst size’) and the initial size of inoculum. This suggests that more productive viruses are better colonizers (in ecology ‘r strategy’). This ‘colonization’ strategy comes at a cost for the virus because infecting the basal layer of the epithelium triggers the immune response. Thus, this strategy colonizes more sites but exploits them less optimally. Another feature that was mediated through the immune response feedback was that mucus trapping delayed the peak of the infection (i.e. the decreased progeny of bacteria and viruses meant less antigen and thus slower immunity detection).

The effect of stage-structure on infection dynamics can be interpreted in light of earlier results from ecology or epidemiology. For instance, in epidemiology, it is known that the more a general population of infected host is subdivided into classes, the more rapid the growth rate of the epidemic is and the shorter it lasts [38]. Our model bears even more parallels with age structured models in epidemiology where the age groups of the host population are explicitly considered. In many of these models, children tend to be key to the spread of epidemics [38], a result that echoes the bottom-up effects we identify. However, the driving forces in the two models are different: in our model it is due to the fact that basal cells are the ones replicating, whereas in epidemiology it is usually driven by longer lasting acquired immunity at higher ages.

### Perspectives

Spatial structure is a natural extension of our model that could be investigated further. Here, the different cell populations partly capture the vertical structure. A specific consequence of not including space is the that the immune system effects are less homogeneous than we have modeled, given that more immune cells are present in the lower layers. The assumption of well-mixed populations holds best when the model represents a portion of the squamous epithelium (rather than say, the whole cervix) because the larger the basal layer, the less applicable the homogeneous mixing assumption becomes. In the case of patchy infections like HPV, a metapopulation modeling approach may be more appropriate (e.g. [24, 60]) or a full spatial model [23]. We chose not to include space since much of the experimental data available on these systems is not spatial. Instead most are cell population counts from immunofluorescence or flow cytometry techniques. However, depending on the experimental setup and questions, space most certainly could be considered in future studies.

Introducing stochastic aspects in epithelial dynamics have recently refueled the discussion on the determinants of HPV clearance [25]. In general, considering stochastic dynamics could matter most when pathogen population approach low-levels (i.e. very few infected cells or small loads). For instance, our finding that mucus trapping can delay the peak and the duration of infections could interact with stochasticity. This is similarly true for infections started with a small inoculum, very rapid abrasion closure, and rapid repair with small inoculum. These processes keep the pathogen populations sizes down and thus, as seen in ecological systems, stochasticity should play a larger role in extinction. As for the spatial structure, it is important to stress that there often is little data on the initial stage of the infections, when the pathogen is rare.

Many previous works have used simplified descriptions of the immune response in a similar fashion as we have chosen to model here [17, 29]. Models with simplified immunity usually ask conceptual questions and/or are used to infer parameter values from data with few measured cell types (e.g. only counting CD8^+^ T and CD4^+^ T cells). Future work interested in specific questions that are immune related, for instance comparing the respective roles of innate and adaptive immunity in clearance or, with more data, could benefit from a more detailed descriptions of immune effectors. In particular, populations of cytokines are interesting as they are important in the epithelium’s role in innate immunity [54].

Our model does not attempt to capture the progression stages that HPVs can cause in persisting infections. To appropriately model these changes would require several adjustments, including that cell proliferation of infected cells and probabilities of symmetric divisions become time variant. Indeed, our model can be adapted to study other oncoviruses that infect the epithelium, where future studies can consider the contexts of immune evasion and cellular transformation driven by oncogenes [42]. In addition, there is increased interest in how epithelial cell dynamics (e.g. cell competition, mechanisms to maintain homeostasis and repair) interact with our knowledge of how tumour viruses alter cellular programing, in particular changing balanced cell fate ratios, skewing squamous differentiation toward a proliferative phenotype [61]. New modeling methods will require possible evolutionary approaches of cell phenotypes emerging over time.

In many ways, the simultaneous infection of a host by different pathogen strains or even species is the rule rather than the exception [62]. Of particular interest is how different pathogens or strains interact inside a host and how this affects the course of the infection.

For instance, HPV infections are often of multiple HPV types and as lesions progress to cancer there is clonal-selection, usually leading to a single type as the main driver of the tumour [63]. One straight-forward extension of this model would be to investigate coinfections between pathogens with similar cell tropisms (e.g. Chlamydia and EBV) or pathogens that differ in their life-cycles. Our model could consider both infections at once or be adapted to study organotypic models that include multiple pathogen infections (e.g. EBV and HPV coinfecting the same tissues and cells [64]) or the effects of the resident microbiota.

Finally, opening a dialogue between mathematical modeling and experimental data generates new hypotheses to test. One of the clearest illustrations of this is our result that burst size differences appear as the most parsimonious explanation to explain symptom differences between wart-causing and lesion-causing HPV infections. Technological improvements in clinical and experimental techniques also allow us to test more subtle predictions. Testing hypotheses generated by the model will allow us to move forward by validating the model assumptions that are consistent with the data and rejecting the others. This will allow us to increase the model complexity and test more elaborate predictions. We hope to inspire experimental studies on infections of stratified epithelia to focus more on dynamics and time series approaches (including mathematics) to better understand these varied and broadly impacting pathogens.

## MATERIALS AND METHODS

### Data

Organotypic culture growing techniques used here have already been described in detail elsewhere [65, 66]. Original experiments were performed to obtain time series data with sufficient replicates for model fitting. Three independent experiments were performed, with rafts harvested at one-week intervals (0, 1, 2, and 3 weeks) starting the day after lifting them to an air-liquid interface. From a total of 12 formalin-fixed, paraffin-embedded (FFPE) rafts, 48 tissue slices were imaged using fluorescence microscopy (DAPI staining for cell nuclei) and resulted in 3 Fields of View (FOV) per slice (*n* = 144). Counts in each FOV were done semi-automatized using ImageJ cell counting software.

### Epithelial model

The uninfected epithelial model consists of 4 cell populations of distinct phenotypes to capture epithelial structure (Figure 1): basal cells (assumed to have a constant population size, *U_b_* = *N_b_*, as cells that move up are replaced immediately), parabasal cells (with population size *U_p_*), differentiated cells of the mid and upper layers (with population size *U_d_*) and of the surface layer (with population size *U_s_*). Since we are interested in cervicovaginal infections which infect non-keratinized squamous epithelia, we assume the top layer of keratinocytes are close to death and that are shed from the surface as they die. The cell population dynamics are captured by three ordinary differential equations:

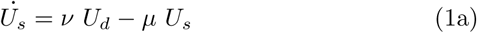

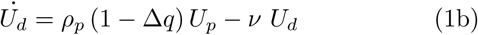

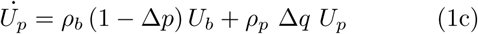

Dots above the variables indicate time derivatives. Basal cells proliferate at a rate *ρ_b_*, giving rise to parabasal cells which in turn proliferate at a rate *ρ_p_*, while entering the mid and upper layers of the squamous column (equation 1). These cells are differentiated and migrate up to the surface layer at a rate *ν* (equation 1b). Mature keratinocytes die at the surface of the epithelium at a rate *μ* (equation 1a). There is an overlap between cell phenotype and spatial structure since an epithelial cell moves up the stages as it ages (Figure 1).

When modeling stem cell divisions, we follow earlier studies [25, 67] and introduce the probability that a basal cell division is symmetric and gives rise to two basal cells (*p*_1_) and the probability that gives birth to two parabasal daughter cells (*p*_2_). Note that *q*_1_ and *q*_2_ are the parabasal equivalent terms (see Figure 1). For simplicity, we only follow the difference of one type of symmetric division (Δ_*p*_ = *p*_1_ − *p*_2_ and Δ_*q*_ = *q*_1_ − *q*_2_ in equations 1b and 1c). We chose to not include the stochastic nature of these divisions, as it has been considered previously [25, 67], and we were interested in understanding deterministic behaviours of the system, such as active repair or active changes to cell ratios. All the variables and parameters used are summarized in Figure 1 and Table I. Finally, the model is sufficiently general that it can represent different kinds of stratified epithelia, including keratinized and non-keratinized squamous epithelia.

To calibrate parameters (Table I), we initially relied on a study from 1970 that used *in vivo* autoradiography techniques to calculate the mean cell cycle time for epithelial cells in cervical and vaginal tissues [45]. They found that basal cells have a relatively slow cycle of approximately 33 days and that 1.14% of these cells are synthesizing DNA at a given time point. Parabasal cells have a much shorter cell cycle (2.6 days) and 14.25% of these cells are synthesizing DNA. Differentiated cells do not divide and have a life expectancy of 4 days (Table I). A detailed analytical analysis of this uninfected model can be found in the Supporting Information.

For fitting raft cell culture data, where the tissue is grown up from a single layer of basal cells, we used a variation of our model. The main difference was assuming the basal layer was not constant but rather followed this equation:
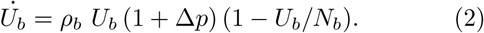

Here we assume the basal layer (cells that are touching the basal lamina) are growing until they reach a maximum capacity, *N_b_*. There are three other changes from the previous model: *U_s_* now represents the surface cells that are keratinized, and since parabasal cells and suprabasal cells counted in the experiment cannot be distinguished we summed these two variables for the fitting. We also assumed *μ* = 0 since the cells are washed away by mucus which is a process not present in our experimental setup.

### Infection models

Modeling infections requires adding populations of infected cells and free-forms of the pathogens. HPV infects basal cells (*I_b_*), which then move up the epithelium column (*I_p_*, *I_d_* and *I_s_*) and release virions (*V*) when dying at the surface (Figure 5A). Similarly, Chlamydia is an obligate intracellular bacteria with an extracellular stage, the elementary bodies (EB). Our model builds on existing models [18, 20], with several adjustments. In particular, to capture the structure of the epithelium, the EBs are divided into an upper (*E_u_*) and a lower (*E_l_*) population, which migrate up and down the epithelium. The resulting life-cycle of Chlamydia is in Figure 5B, and all equations are in the Supporting Information.

**FIG. 5.**
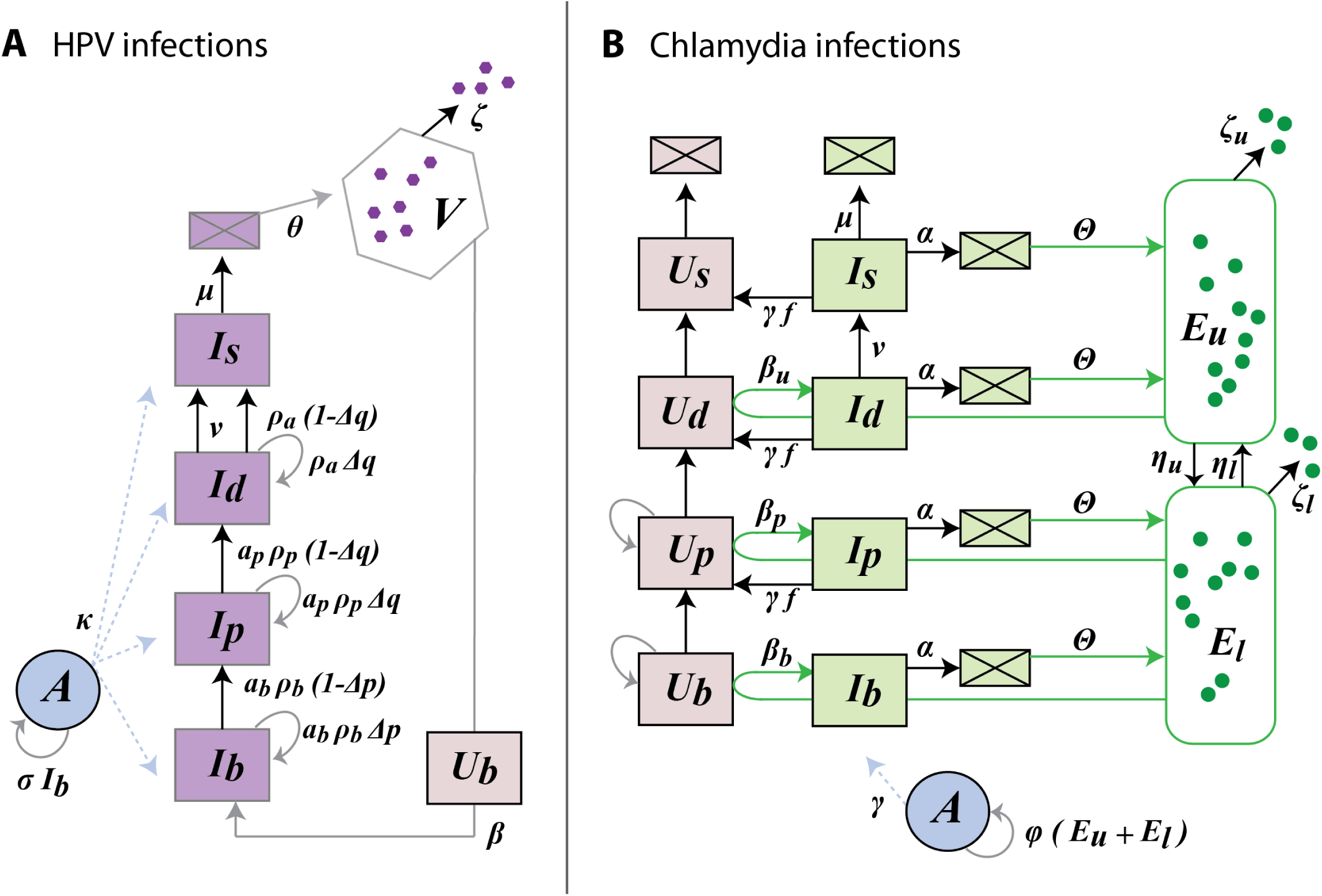
Flow diagram of the infection models for HPV (A) and Chlamydia (B). HPV is non-lytic and its life-cycle follows that of epithelial cells. In the case of *C. trachomatis*, the elementary bodies, EBs, can be found either in the top layers, *E_u_*, or in the lower layers, *E_l_*. EBs carry the migration of the infection between layers, *η*. Finally, the host immune response, *A*, is activated by infected basal cells in the case of HPV and all EBs in the case of *C. trachomatis*. Note that for LR-HPV infections *ρ_a_* = 0 and *α_b_* = *α_p_* = 1.

For both infections, following earlier studies (e.g. [29]), we simplify the immune response to a single population of CD8^+^ cells (Figure 5). As shown in the Results, this simple formalism is sufficient to capture important feedbacks and investigate the effects of parameters on infection duration. Investigating more subtle immunity dynamics or data fitting would call for a more detailed description of the immune response, which is beyond the scope of this work.

Infection dynamics were simulated in Mathematica [68] using NDSolve (with methods ‘BDF’ or ‘StiffnessSwitching’) for numerical integration.

### Parameter values and sensitivity analyses

Nearly all the parameter values could be set using data from the literature, which mostly lay in narrow ranges (Table I and Table S1). Parameters for which we had little information were either kept free or calibrated. For instance we used Δ_*q*_ to scale all equilibrium population sizes (see the Results).

To test the robustness of our results, we performed uncertainty and sensitivity analyses using Latin Hypercube Sampling and Partial Rank Correlation Coefficients (PRCC) via the pse package in R [69], which is popular for disease models [70], and numerical integration was done using `deSolve` package. We generated 1,000 parameter sets by Latin Hypercube sampling from uniformly distributed parameter values within a specified biologically realistic range. PRCCs were calculated between the rank-transformed samples and the resulting output matrix of the response variables (e.g. duration of infection, maximum pathogen load). 100 bootstraps were performed to generate 95% confidence intervals. The magnitude of the PRCCs determines the effect strength of a given parameter on a specific response variable (0 for no effect and 1 for very strong) and the sign indicates whether the response grows or shrinks with increasing the parameter value.

Monotonicity for each parameter was checked for each response variable, and the parameter ranges were shortened when monotonicity was not obeyed. This was not common and was usually for values very close to zero.

### Parameter estimation from experimental data

We inferred parameter values from the data we collected over 3 weeks from a 3D raft culture of NIKS cells. Note that cells attached to the basal membrane were considered basal and those above them were counted as suprabasal cells. This was done (rather than use differentiation markers) in order to differentiate between true basal cells and parabasals and to estimate a carrying capacity, *N_b_*. Model parameters were inferred using maximum likelihood estimation and trajectory matching, assuming measurement error follows a Poisson distribution. Fitting and model predictions were performed in R software [71], using packages `bbmle` [72], `deSolve` [73], and `pomp` [74].

## ACKNOWLEDGMENTS

We would like to thank Kathlyn Alexander for assistance with the raft culture experiments and also to Drs. Ignacio G. Bravo, Jessie L. Abbate and Jérémie Guedj for helpful discussions. We also thank Dr. Paul Lambert (McArdle Laboratory for Cancer Research, University of Wisconsin) for providing the NIKS cell line.

